# Adjustable Compliance Footwear Technology to Investigate Gait Adaptation

**DOI:** 10.1101/2025.03.31.645746

**Authors:** Mark Price, Calder Robbins, Sarah Szemethy, Banu Abdikadirova, Gina Olson, Wouter Hoogkamer, Meghan E Huber

## Abstract

Kinetic asymmetries in gait caused by neurological impairments are a major factor in mobility deficiency. Gait perturbation training has shown promise for rehabilitating neurological gait asymmetry, but the transfer of benefits from treadmill training to overground walking is limited. To enable kinetic gait perturbation training on treadmills and overground, we developed novel adjustable compliance footwear technology. The system consists of a wearable pneumatic actuation module which controls the pressure of custom air pockets embedded in a soft lattice midsole, and is capable of independently and wirelessly adjusting the midsole compliance under each foot from 300 kN/m and 60% energy absorption to 18 kN/m and 75% energy absorption in under 4 seconds. Hindfoot and forefoot sole deflection data are measured via Hall effect sensors and paired magnets characterized to 1.7 mm accuracy. To demonstrate the experimental capability of the system, we successfully triggered multiple compliance changes during overground walking and recorded ground reaction forces and embedded sensor values during a prolonged asymmetric compliance walking bout on a treadmill. Future work will include a full experimental study of gait adaptation to asymmetric footwear compliance perturbations.

## I. INTRODUCTION

Impaired locomotion and balance are a highly prevalent cause of mobility loss, decline in independence, and the development of debilitating comorbidities. Stroke, for example, is a leading cause of long-term disability in the US and leads to gait impairments in 80% of cases [1]. A major disabling factor in neurologically impaired gait is the loss of ability to provide symmetrical weight bearing and propulsion forces between limbs. Deficiencies in paretic propulsion, or the percentage of propulsive impulse generated from the paretic side limb, are strongly correlated with hemiparetic severity and lower gait performance in general [3]. Furthermore, walking with weight-bearing asymmetries can eventually lead to osteoarthritis if uncorrected [4], [5]. Symmetry in kinetic measures of gait is therefore a crucial target for neurorehabilitation.

Eliciting adaptations via asymmetric gait perturbations has emerged as a promising technique for restoring symmetry in neurorehabilitation. The technique of running treadmill belts at different speeds under each foot of a walker (i.e., split-belt treadmill training) is capable of eliciting changes in spatial measures such as step length, both in the form of immediate after-effects lasting up to a few minutes after first exposure [6], [7] to more permanent changes that remain months after repetitive training [8]. Similar approaches have more recently been developed to asymmetrically vary the vertical surface stiffness under each foot on a treadmill with the aim of eliciting changes in muscle coordination and ground reaction forces [9], [10]. In these approaches, encouragement of more symmetrical weight bearing is hypothesized to stimulate active motor signals from the central nervous system to the paretic side limb, which may elicit desirable outcomes such as increased usage of the paretic leg, stimulated neuromotor adaptation, and restoration of function through neuroplasticity [10].

Despite the promise of these approaches, treadmill-based neuromotor adaptation paradigms do not fully transfer their effects to overground walking. This gap is not fully understood, but it has been proposed that changes in perceptual context between learning and application limit the transfer of learned skills [11], including split-belt walking behaviors [7], [12], [13]. Furthermore, a treadmill constrains the adaptation behavior to maintain a near-constant walking speed and limited medio-lateral motion to remain on the belts. Gait asymmetries in people with chronic stroke [14], [15] or other causes (i.e., amputation [16], [17]) present differently on treadmills and overground, despite otherwise matched conditions. Limited transfer of training to overground contexts is a critical limitation; the benefits of training do not fully translate to day-to-day life.

We propose that a tool to deliver adjustable foot-ground compliance gait training overground would remove this training context barrier. In this paper, we present and evaluate a design for such a tool. The contributions of this work are: (1) Development of an overground tool will allow training to occur in naturalistic contexts and will not constrain the walking speed of the user, allowing them to adapt as they would in everyday scenarios and potentially increasing the training benefits. (2) An overground tool will require a design that is wearable, portable, and untethered, which will more easily translate into an accessible take-home rehabilitation tool for individuals with impaired gait, requiring substantially less material and infrastructural investment than the installation of a specialized treadmill. (3) A wearable adjustable stiffness tool that functions overground will naturally also function on treadmills, allowing controlled study of the effect of treadmill vs. overground adaptation to stiffness perturbations.

To provide these capabilities, we designed portable robotic footwear capable of changing foot-ground compliance in real time to deliver perturbations during overground walking. In this paper, we present the design of the system, characterize its performance as a variable stiffness device, validate its function as experimental hardware with two human participants, and discuss the findings of our evaluations.

## II. DESIGN

The portable robotic footwear system comprises two subassemblies: (1) custom soft shoe soles with embedded pneumatic variable stiffness pockets, and (2) a pressure control system mounted on a wearable harness. These assemblies were designed to meet criteria determined by existing variable stiffness treadmill technology and the demands of performing overground experiments.

### A. Design Specifications

As a mobile alternative to adjustable stiffness treadmill technology, the design of the adjustable compliance shoes is centered on providing similar capabilities to adjustable surface stiffness treadmills [9], [10]. The shoes are therefore designed to: (1) independently change compliance under each foot, allowing for asymmetric ground compliance perturbations, (2) achieve a compressive stiffness range of approximately 15–300 kN/m, (3) provide substantial vertical foot deflection (i.e., greater than 1 cm), (4) change compliance via operator trigger during gait without requiring the user to stop, and (5) change compliance across the full range within 5 strides. This final specification is relaxed from the response time of the treadmills (within one stride) to account for the practical limitations of a fully wearable system.

To prevent the system from overly disrupting gait in the stiff baseline condition, there are additional limits placed on the design: (1) The trunk-suspended mass is limited to 5 kg (actual mass: 5.1 kg) and the mass of each shoe attachment is limited to 1 kg (actual: 0.6 kg), (2) the total stack height is limited to 10 cm (actual: 6.2 cm), which limits the maximum achievable vertical deflection.

Adjustable compliance mechanisms for footwear exist in the literature. Magnetorheological actuators mounted in an array in a shoe midsole have been used to selectively change the compliance beneath specific high-pressure areas of a user’s foot to prevent the development of pressure ulcers for diabetic users [18]. In another example, vacuum-driven layer jamming has been used to increase the stiffness of a passively compliant sole [19]. In both of these cases, the design enables rapid compliance change of a low-deflection material, primarily changing pressure distribution and the sensory experience at the sole of the foot, rather than large deflections at the foot-ground interface as caused by adjustable stiffness treadmills. Inflatable shoe soles did achieve changes in compliance with associated large deflections [20], though the foot-ground interface geometry was substantially changed by the inflation state of the device. Shoe height asymmetry has been observed to cause reactive gait changes [21], which suggests that shoe geometry effects may interact with effects caused by our intended compliance perturbation. Therefore, we pursued a design that uses pneumatic pressure to control compliance of a high-deflection interface with geometry constraints that maintain the shape and height of the interface across the pressure range while unloaded.

### B. Pneumatic Shoe Soles

The adjustable compliance sole consists of custom 3D printed air pockets at the forefoot and hindfoot supported by a passive lattice structure (Fig. 1). The air pockets consist of two flat 3mm panels connected by a thin side wall membrane and thin flexible ribs that attach to the interior surface of each panel. These ribs act as tension members to enforce the parallel geometry and maximum distance between the top and bottom surfaces, and their buckling behavior defines the minimum passive unpressurized stiffness of the air pocket.

**Fig. 1.**
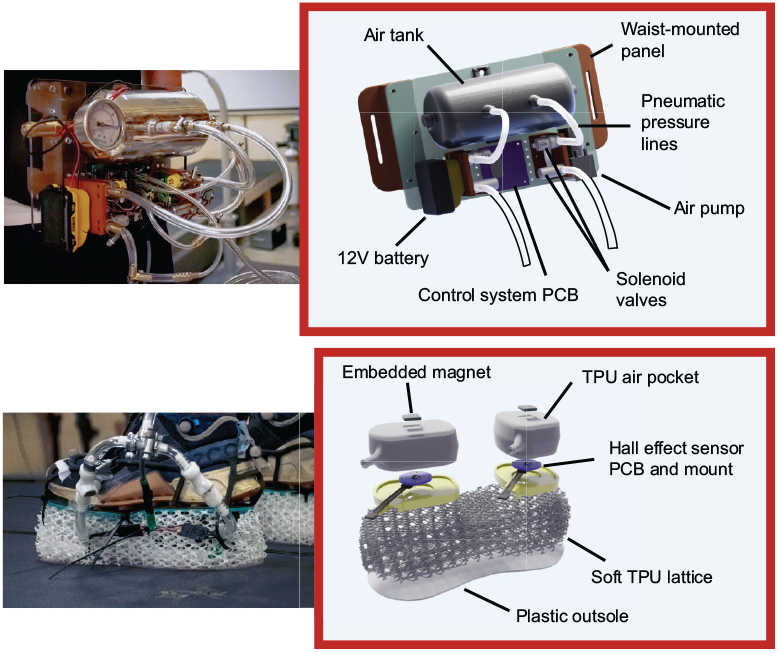
Full robotic shoe assembly with selected labeled components.

Custom printed circuit boards with Hall effect sensors (TMAG5273, Texas Instruments) are mounted under each air pocket, and a 19×19×1.6 mm square axially magnetized magnet is embedded into the top surface of the pocket centered above the corresponding sensor. Magnetic flux density measurements correspond with proximity of the magnet to the sensor, and therefore provide an estimate of deflection at fore- and hindfoot. Both air pockets and sensors are fitted against a rigid plate to restrict free movement of the components during locomotion.

The air pockets and sensor assemblies are encased in a soft lattice structure to protect the assemblies from abrasion, fix their position relative to the foot, and provide a stable, continuous interface between the foot and the ground. The air pockets were designed in Solidworks 3D CAD software and the surrounding lattice was designed with geometry nodes in Blender 3D design software. These structures were fabricated with a fused deposition modeling process (Bambu X1-Carbon, Bambu Labs) extruding flexible TPU filament (Overture High Speed TPU, Shore 95A).

The full adjustable compliance sole assembly is secured to a shoe balancer (EVENup, OPED Medical Inc.) with ratcheting cable ties threaded through the interstitial sites of the lattice. Users may strap the assembly over their daily-use shoes. When fully pressurized, the assembly behaves like a 5cm platform shoe. Stiffness decreases with pressure to the minimum condition where air is allowed to flow freely in and out of the pockets.

### C. Wearable Pressure Control System

Pressure inside the air pockets is controlled with a pneumatic system comprising a pump, air tank, and flow control solenoids (Fig. 1). The system is configured with two independent pneumatic lines, allowing either independent control of two shoe soles with coupled fore- and hindfoot air pockets (as implemented in this paper), or independent control of a forefoot and hindfoot on a single shoe.

#### 1) Hardware

Each pneumatic output is regulated through the control of a flow-control solenoid (4916K7, McMaster) and a cutoff solenoid (5001T35, McMaster). Pressure in both outputs and in the air tank are sensed via analog gauge pressure sensors (SSC series, Honeywell) and are regulated via control of the solenoids and the tank pump, respectively. The air tank is rated for a maximum of 180 psi, but a safety release valve limits the stored air pressure to 30 psi. Control states may be toggled remotely via RF receiver by an operator using paired keyfob transmitter (RF controller, Parallax Inc.). Pressure readings and Hall effect sensor readings are routed through a microcontroller (Arduino Pro Mini) sampling at 100 Hz, which controls the air pump and pressure control solenoids by driving MOSFET power transistors to regulate tank pressure and pressure in both outputs. Signals and information handled by the microcontroller are logged locally on a microSD card. The system is powered by a 12V lithium ion battery.

#### 2) Supply Tank Pressure Control

Pressure in the air tank is maintained between 20-25 psi to allow rapid adjustments in output pressure. This pressure range is controlled by switching on the air pump when pressure falls below the lower range and switching it off when pressure reaches the upper range.

#### 3) Shoe Pressure Control

Output pressure is controlled by switching between three flow control states: pressurize, exhaust, and occlude (Fig. 2). These states are achieved through energizing the two solenoids in different combinations. When sensed pressure exceeds or falls below desired pressure by a margin of +/-0.5 psi, the controller commands the exhaust or pressurize state, respectively. When pressure is within the acceptable range, the occlude state is maintained, trapping the air inside the air pocket. While linear feedback control driving pressure error to zero (i.e. PID control) is possible with this system, it requires constant active solenoid switching, producing a noticeable and continuous disturbance in the compliance under each foot, therefore we prioritized a controller that ignores miniscule adjustments in favor of presenting a constant set of interaction dynamics.

**Fig. 2.**
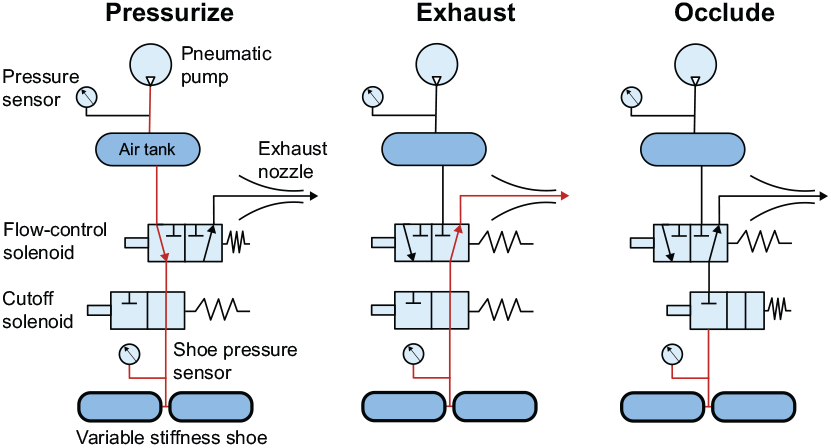
Pneumatic flow control state diagram with airflow highlighted for the flow control states achieved by switching two solenoids in combination. The flow-control solenoid is a 3-way, 2-state valve that either connects the pneumatic shoe to the pressure tank or to the exhaust nozzle. The cutoff solenoid simply connects or disconnects the shoe to the rest of the pneumatic system. This scheme was implemented for both shoes independently.

#### 4) Compliance Control for Locomotion

The adjustable compliance shoes are designed to achieve controllable compliance behavior by trapping a known mass of air (measured as pressure) inside the midsoles. This trapped air combined with the mechanical properties of the midsole structure combine in parallel to form an air spring with characterizable passive compliance properties. Crucially, air pressure is proportional to mass only when the other variable parameters in the ideal gas law are held constant: temperature and volume. Air pocket volume will change under load by design—therefore the passive stiffness is characterized by the air pressure when the shoes are unloaded. The top level control maintains a trapped mass of air associated with the unloaded air pressure for a given compliance condition by allowing air to escape the air pockets only when the target pressure decreases. When the new target is met, instances of actual pressure exceeding the target pressure will trigger the occluded state. This occludes the air pockets during standing or the stance phase of walking and allows adjustments back up to the target unloaded pressure during swing.

## III. EVALUATION

To characterize the mechanical properties of the footwear, evaluate the performance specifications of the mechatronic system, and evaluate the ability of the device to fulfill its role as fully wearable experimental hardware for locomotion experiments, we performed a series of tests. The evaluation tests were designed to: 1) quantify the actual compliance characteristics, the stiffness range, and the speed of maximal stiffness change; 2) quantify the accuracy of measurements calculated from embedded sensor values; 3) evaluate the high level control performance during locomotion.

### A. Adjustable Compliance Evaluation

#### 1) Compliance Characterization

We characterized the compliance of the footwear by compressing a full sole assembly between two rigid plates in a material testing machine (Instron 68TM-10). Three loading conditions were applied: distributed load across the entire sole via a large flat plate, concentrated load above the forefoot pocket via a flat cylinder 63 mm in diameter, and an identical concentrated load above the hindfoot pocket. For the full-shoe load, the machine compressed the shoe at a rate of 5 mm/s until it reached a threshold of 1200 N from a preload of 20 N, paused for 10 s, then decreased back to the 20N preload at the same rate. The concentrated loads were applied to reach 800 N with the same profile. This process was repeated 3 times for 7 pressure conditions: exhaust mode (air is allowed to freely escape), and occluded 0, 3, 6, 9, 12, and 15 psi. Force vs. displacement curves were calculated from the load cell and cross-head position outputs of the machine.

Force vs. displacement curves are shown in Fig. 3. As may be expected from air springs and soft materials, the stiffness profile is nonlinear. To represent the mechanical behavior of the system, we split the positive loading response into low load and high load sections at a threshold of half of the maximum applied load. We applied a linear regression to all data points within each section across the three cycles per pressure condition to estimate a linear spring coefficient for the low-load and high-load response separately. Because we cannot identify a linear damping ratio for such nonlinear system dynamics, we calculated the energy input to the shoe and the net energy dissipate by the shoe during each cycle by integrating the applied load with respect to displacement during loading, and during and unloading (i.e., the area within the hysteresis curve), respectively. We reported the average compressive work and net work per loading cycle for each pressure condition in Fig. 3.

**Fig. 3.**
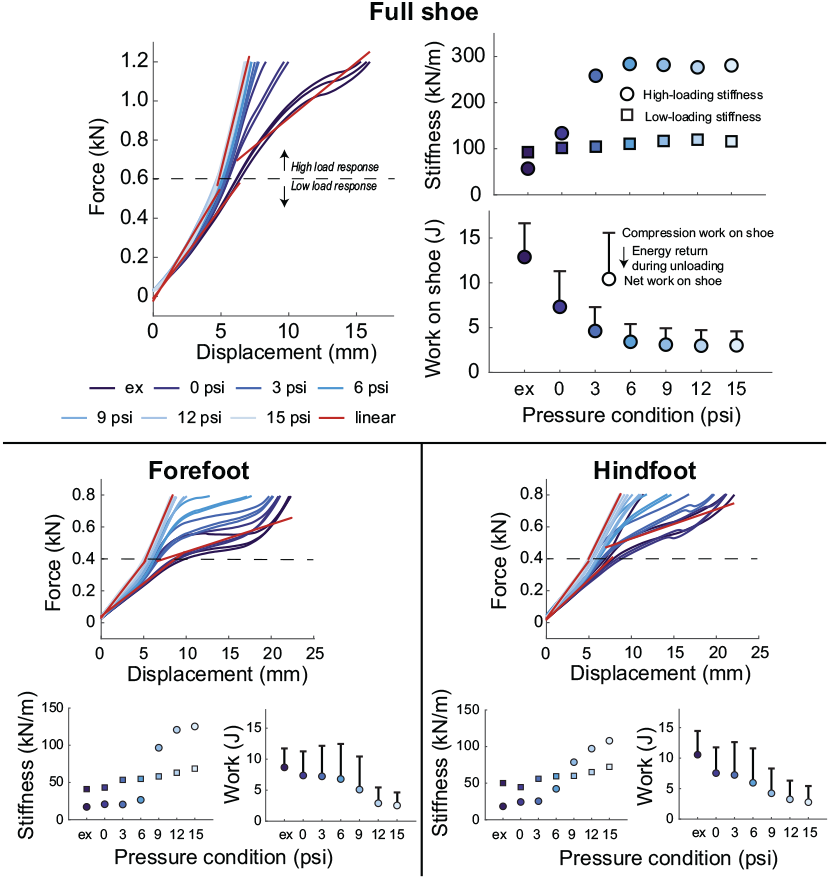
Force vs. displacement curves for 7 pressure control states for an evenly distributed full shoe load, a concentrated forefoot load, and a concentrated hindfoot load. Low- and high-load thresholds for characterizing stiffness are indicated with a dashed line. Linear regression lines for low- and high-load ranges are plotted for the the minimum (exhaust state (ex)) and maximum (15 psi) stiffness conditions for visual reference to Table I. The slope of each regression line is plotted as a stiffness value. Positive work done on the shoe as well as the net energy dissipated by the shoe per loading cycle are indicated for each pressure condition.

Linear regression for the low load force-displacement relationship achieved *R*^2^ *>* 0.94 for all conditions. Both stiffness coefficient magnitude and *R*^2^ values for the high load response were more variable, and are reported in Table I. The stiffness when distributing load over the full shoe surface is overall higher than when concentrating load over one pocket, ranging from approximately 57–280 kN/m at high loads. This stiffness plateaus above 6 psi for the full shoe, but at ∼12 psi for concentrated loads.

**Table I.** Robotic shoe stiffness and energy dissipation

| Loading | Measure | exhaust | 0 psi | 3 psi | 6 psi | 9 psi | 12 psi | 15 psi |
| --- | --- | --- | --- | --- | --- | --- | --- | --- |
| Full shoe | High-load K (kN/m) | 56.8 | 133.6 | 258.3 | 283.8 | 281.9 | 276.6 | 280.7 |
| | $R^2$ | 0.92 | 0.88 | 0.99 | 0.99 | 0.99 | 0.99 | 0.96 |
|  | Energy lost/Energy in | 77.4% | 64.8% | 63.5% | 63.3% | 62.9% | 63.1% | 65.9% |
| Forefoot | High-load K (kN/m) | 17.2 | 20.7 | 20.4 | 26.6 | 96.7 | 120.8 | 125.3 |
| | $R^2$ | 0.57 | 0.72 | 0.84 | 0.70 | 0.94 | 0.98 | 0.99 |
|  | Energy lost/Energy in | 74.0% | 65.5% | 59.5% | 54.6% | 48.8% | 53.1% | 54.7% |
| Hindfoot | High-load K (kN/m) | 18.3 | 24.3 | 25.4 | 42.2 | 78.7 | 97.1 | 107.9 |
| | $R^2$ | 0.58 | 0.85 | 0.85 | 0.83 | 0.95 | 0.98 | 0.99 |
|  | Energy lost/Energy in | 73.1% | 64.1% | 57.3% | 51.3% | 51.0% | 51.4% | 51.1% |

All conditions are highly dissipative, with less than half of the mechanical energy input to the shoe being returned during unloading for all conditions, with notably more energy dissipation occurring during the exhaust condition (over 70% energy dissipation for all loading conditions). The percentage of energy dissipated by the shoe relative to the work done to it during compression is also reported in Table I.

#### 2) Stiffness Control Step Response

We tested the time required to achieve a full bandwidth compliance change for one shoe with and without a bodyweight load (792 N) concentrated on that shoe. We measured the 90% rise time from a 12.5 psi pressure command from the exhaust state, and an exhaust state command from 12.5 psi, averaged across 5 trials for each state change and each loading condition (total of 20 trials). For the unloaded condition, the rise time was 3.41 s and the fall time was 3.80 s. For the loaded condition, the rise time was 3.63 s and the fall time was 4.14 s. The step response profile is illustrated in Fig. 4.

**Fig. 4.**
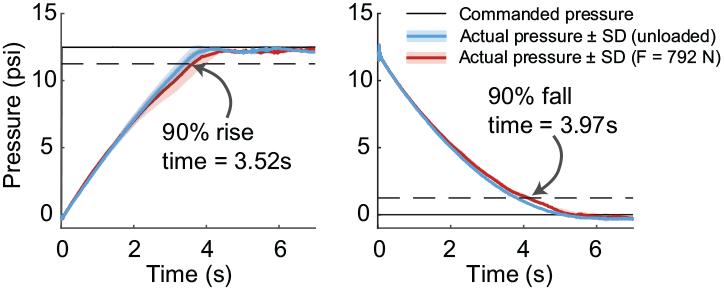
Step response for maximal stiffness change (i.e., transition from exhaust condition to 12 psi). Rise and fall times reflect the average between loaded and unloaded conditions—both times increased by ∼ 300 ms with the standing weight of a 80.7 kg person applied. Traces reflect the average and standard deviation of 5 trials across loaded and unloaded conditions.

### B. Embedded Sensor Performance

#### 1) Deflection model and evaluation

We quantified the relationship between vertical magnet deflection and magnetic flux density by suspending a magnet from the material testing machine crosshead centered above the embedded Hall effect sensor, then moving it from 10.02 mm to 51.66 mm above the sensor. We continuously recorded data at 100 Hz through this motion, resulting in about 7,000 samples. We then fit an inverse cube root model to the magnet distance vs. flux density curve (*R*^2^ = 0.999, RMSE = 0.45 mm, Fig. 5A).

**Fig. 5.**
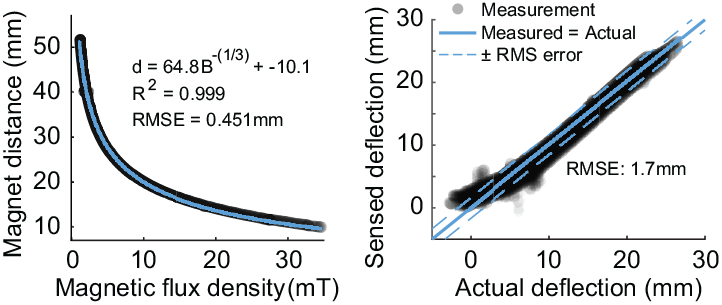
Hall effect sensor displacement model fitting and evaluation. A: Model fitting with ideal vertical displacement conditions. B: Sensed deflection from the Hall effect model vs. actual deflection measured by the material testing machine under actual compression loads.

We evaluated this model by calculating vertical magnet deflection from flux density measurements recorded during the stiffness characterization experiments for concentrated load above each air pocket, and compared these values to the corresponding crosshead position, combining to ∼125,000 samples. The model estimated vertical deflection with RMS error of 1.66 mm (Fig. 5B).

### C. Adjustable Compliance Locomotion

To demonstrate the capability of the device to achieve changes in foot-ground stiffness during walking, we performed two tests with human pilots: (1) Overground locomotion over varied terrain while foot-ground stiffness was adjusted across the range of available values, and (2) a gait adaptation protocol to an asymmetric ground stiffness perturbation on a rigid instrumented treadmill. Approval for these tests was obtained from the host university’s Institutional Review Board (Protocol #3638).

#### 1) Overground locomotion proof-of-concept

A volunteer pilot (64.5 kg) wore the robotic shoe system and navigated a hallway containing a 90 degree turn, a short set of 3 stairs, and a ∼ 8% incline at a self-selected speed. An operator followed behind and wirelessly triggered changes to an arbitrary set of pressure control targets within the characterized range. All compliance transitions were successfully achieved during walking, as indicated by the corresponding increases and decreases in midsole compression (Fig. 6).

**Fig. 6.**
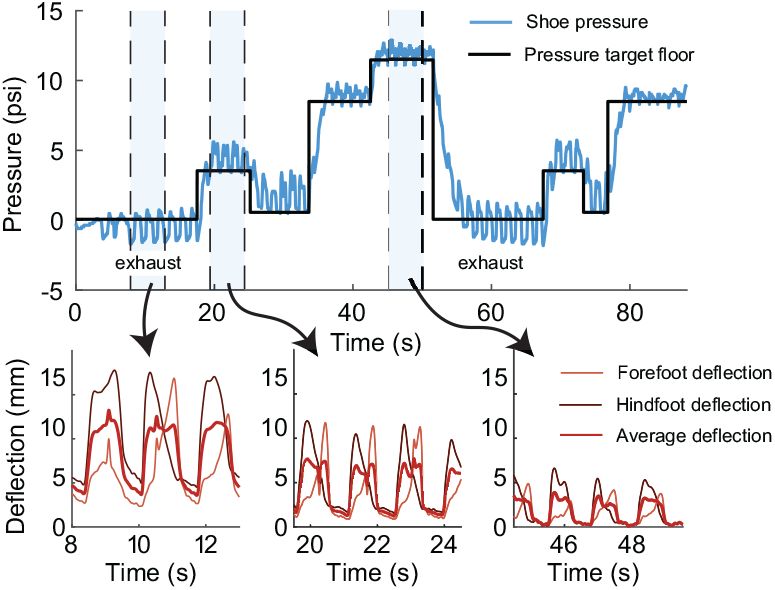
Pressure tracking and resulting shoe deflections during overground walking at a self-selected speed. Note that in the “exhaust” state, sensed pressure is negative during swing as air is displaced out of the system during stance.

#### 2) Asymmetry adaptation on a rigid treadmill

We asymmetrically perturbed the shoe compliance for 10 minutes during 1.0 m/s treadmill walking with one volunteer (66.3 kg) to demonstrate the capability of the footwear to replicate gait adaptation experiments conducted with asymmetry perturbations [6], [7] (Fig. 7). After walking for 5 minutes with both shoes at the 12 psi condition, the exhaust condition was triggered on one side only. Symmetrical pressure states were restored after 10 minutes of exposure. We recorded 3D ground reaction forces from the treadmill (Bertec Fully Instrumented Treadmill) and averaged 10-stride windows immediately before the perturbation, at the end of the perturbation, and four strides after the perturbation (accounting for time to reach the pressure target) to observe gait changes due to the change in compliance. Results from this pilot indicate that the footwear perturbation causes qualitative changes in walking ground reaction forces during and after exposure to footwear compliance asymmetry (Fig 7), though further study with more participants will be required to determine quantitative effects.

**Fig. 7.**
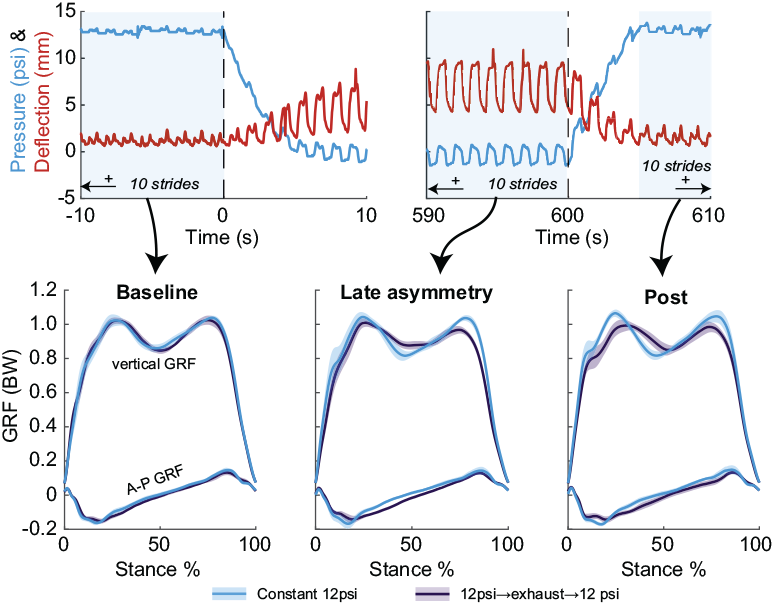
Pilot results from a 10-minute asymmetric ground stiffness perturbation applied by the robotic shoes while walking at 1.0 m/s on an instrumented treadmill. Pressure and deflection values reflect the embedded sensor readings in the shoe that changed state; deflection is the mean of the hindfoot and forefoot sensors. Ground reaction force profiles reflect mean and standard deviation of 10 gait cycles per window.

## IV. DISCUSSION

We designed and built adjustable compliance footwear capable of changing the foot-ground interface compliance independently under each foot during locomotion within a few steps. Preliminary tests with volunteer pilots show promise toward the use of the footwear for locomotion adaptation experiments. Though the evaluation indicates that the novel system is a functional experimental tool, here we compare the results to the closest direct analogue: adjustable stiffness treadmills.

### A. Comparison with adjustable stiffness treadmills

By nature of being an untethered, wearable system, there are unavoidable differences between the footwear and adjustable stiffness treadmills. Unlike with a suspended treadmill platform, the footwear compliance depends on the concentration of load, and are much more compliant for loads concentrated above one of the air pockets (e.g., at the start and end of stance). Additionally, the footwear have highly dissipative and nonlinear loading dynamics.. While adjustable mechanical spring mechanisms with more ideal springlike behavior than our pneumatic system have been implemented in prosthetic feet [22], [23], these mechanisms control bending stiffness, and would require a considerably larger footprint to apply to vertical compression stiffness over the whole foot. There is evidence from optimal control simulations of asymmetric surface compliance walking that damping may have a major role in the resulting gait pattern [24]. While not a perfect match for the simulated system dynamics, the dissipative footwear response may provide an opportunity to investigate these effects experimentally.

Furthermore, the footwear has geometric and power limitations required to make the system practical as an untethered wearable apparatus. We measured over 2 cm of vertical deflection during our tests, compared with the 10 cm travel limit of the latest adjustable stiffness treadmill technology [10] [9]. This is a compromise between perturbation magnitude and the degree to which “unperturbed” walking is affected by wearing the footwear in their maximally stiff control state. The time required to complete a step change in stiffness is greater than the step response time of adjustable stiffness treadmills (*<* 1 s, [9], [10]). This is limited by the flow rate of the solenoid valves—faster state transitions are possible if larger (and therefore heavier) solenoids are used.

## V. CONCLUSION

The adjustable compliance footwear system is a functional experimental tool for studying gait adaptations to changing terrain compliance. Future work will focus on using this hardware to study gait adaptation effects and after-effects both in treadmill walking and overground gait training.

